# Impaired glycolysis in bone underlies age-related bone loss in mice

**DOI:** 10.64898/2026.10.06.738500

**Authors:** Fangfang Song, Tyler Marmo, Won Dong Lee, Jasmin Koehnken Sawall, Chao Song, Xueyang Liao, Xing Ji, Karthi Chellappa, Melanie R McReynolds, Joseph A. Baur, Christopher Thom, Fanxin Long

**Affiliations:** Translational Research Program in Pediatric Orthopedics, The Children’s Hospital of Philadelphia; State Key Laboratory of Oral & Maxillofacial Reconstruction and Regeneration, Key Laboratory of Oral Biomedicine Ministry of Education, Hubei Key Laboratory of Stomatology, School & Hospital of Stomatology, Wuhan University, Wuhan 430079, China; Lewis Sigler Institute for Integrative Genomics, Princeton University, Princeton, United States; Department of Biochemistry, Yonsei University, Seoul, Republic of Korea; Institute for Diabetes, Obesity, and Metabolism and Department of Physiology, Perelman School of Medicine, University of Pennsylvania; Department of Biochemistry and Molecular Biology, The Huck Institute of Life Sciences, Pennsylvania State University, University Park, Pennsylvania, USA; Division of Neonatology, Children’s Hospital of Philadelphia, Philadelphia, PA; Department of Pediatrics, University of Pennsylvania Perelman School of Medicine, Philadelphia, PA; Department of Orthopedic Surgery, University of Pennsylvania, Philadelphia, PA

## Abstract

Bone loss and metabolic dysregulation often coexist in older adults, yet the relationship between age-related changes in bone cell metabolism and bone-forming activity remains unclear. By modeling both conditions in aged mice and using in vivo stable-isotope tracing, we demonstrated that age-related decline in bone formation was coupled with an intrinsic impairment of glycolysis within bone tissue. Impaired glycolysis was associated with reduced protein levels of key glycolysis enzymes, and transcriptomics analyses revealed compensatory upregulation of glycolysis genes in aged bone. Metabolic intervention through long-term caloric restriction prevented age-related trabecular bone loss. Furthermore, osteoblast-specific genetic enhancement of glycolysis—via overexpression of HIF1α or the rate-limiting enzyme PFKFB3—significantly improved osteoblast function and increased bone mass in aged mice. Together, the study identifies glycolytic decline as a cell-autonomous driver of osteoblast deficiency and provides proof of principle that reactivating glycolysis can promote bone accrual in aging.

## Introduction

Age-related osteoporosis represents a significant and growing global health burden, characterized by a progressive decline in bone mass and deterioration of bone microarchitecture, leading to increased fragility and fracture risk (1). Bone is a dynamic organ with mechanical and endocrine functions that undergoes continuous regeneration through a bone remodeling process driven by the bone-resorbing osteoclasts and bone-forming osteoblasts (2). After reaching peak bone mass in the third decade, healthy individuals maintain a steady-state bone mass for approximately three decades through the tight coupling of bone resorption and formation (3). With aging, bone remodeling shifts to a low turn-over state, creating a net bone loss due to an imbalance favoring bone resorption over bone formation (4). Compared to postmenopausal bone loss, the mechanisms underlying age-related bone loss in both women and men are understudied. Previous studies have implicated cellular senescence and increased reactive oxygen species in bone aging, but a full understanding of the age-dependent decline in osteoblast differentiation and function remains elusive (5, 6). Furthermore, while aging is known to cause both metabolic dysregulation and osteoporosis, the specific age-related metabolic changes within osteoblasts and their direct contribution to bone loss are not clear.

The maintenance of bone mass is a highly energy-demanding process that not only responds to but also modulates whole-body metabolism (7). Osteoblasts, as the chief bone-making cells responsible for bone matrix synthesis and mineralization, require substantial amounts of adenosine triphosphate (ATP) and biosynthetic precursors that are extracted from various energy substrates including glucose, fatty acids and amino acids (8–12). The metabolic behavior and substrate utilization of osteoblast lineage cells likely change according to their differentiation stage and biosynthetic demand (13). We and others have shown that bioenergetics shifts towards a greater reliance on aerobic glycolysis with osteoblast differentiation in vitro (14, 15). The predominate fate of glucose toward lactate production in bone was validated by glucose tracing experiments in vivo (14). Moreover, emerging evidence indicates that dysregulation of cellular metabolism in osteoblasts contributes to bone disorders (12). Our recent studies identify glycolytic dysfunction as a critical mechanism causing bone loss in mouse models of type I or type II diabetes (16, 17). Thus, research to date supports an integral role for cellular metabolic pathways in mediating osteoblast differentiation and function.

Here, we demonstrate a significant downregulation of glycolytic activity in the bone tissue of aged mice. Genetically enhancing glycolysis through overexpression of key regulators HIF1α or PFKFB3 is sufficient to stimulate bone formation and ameliorate bone loss in aged mice. These findings highlight that restoring glycolytic metabolism is a novel anabolic strategy that targets the underlying bioenergetic deficiencies in aged bone.

## Results

### Bone loss is associated with low turnover and impaired glucose handling in aged mice

To investigate the mechanisms underlying age-related osteoporosis, we first characterized the skeletal phenotype of young (4-month-old) and aged (23-month-old) male mice. Micro-computed tomography (μCT) analysis of the proximal tibia revealed a dramatic loss of trabecular bone in aged mice compared to their young counterparts (Fig. 1A, upper panel). Quantitative analyses confirmed this observation, showing a significant reduction in bone volume fraction (BV/TV), trabecular number (Tb.N), and trabecular thickness (Tb.Th), accompanied by a marked increase in trabecular separation (Tb.Sp) (Fig. 1B). Furthermore, analysis of the mid-diaphysis demonstrated that cortical bone was also adversely affected by aging, as evidenced by significant decreases in cortical bone area and thickness (Fig. A, lower panel, C). In a separate cohort of young (5-month-old) versus old (20-month-old) mice, serum biochemistry showed that both PINP, a marker of bone formation, and CTX-1, a marker of bone resorption, were downregulated in the old group, indicating reduced bone turnover (Fig. 1D, E). Thus, aging in mice causes significant bone loss driven by diminished bone formation.

**Figure 1.**
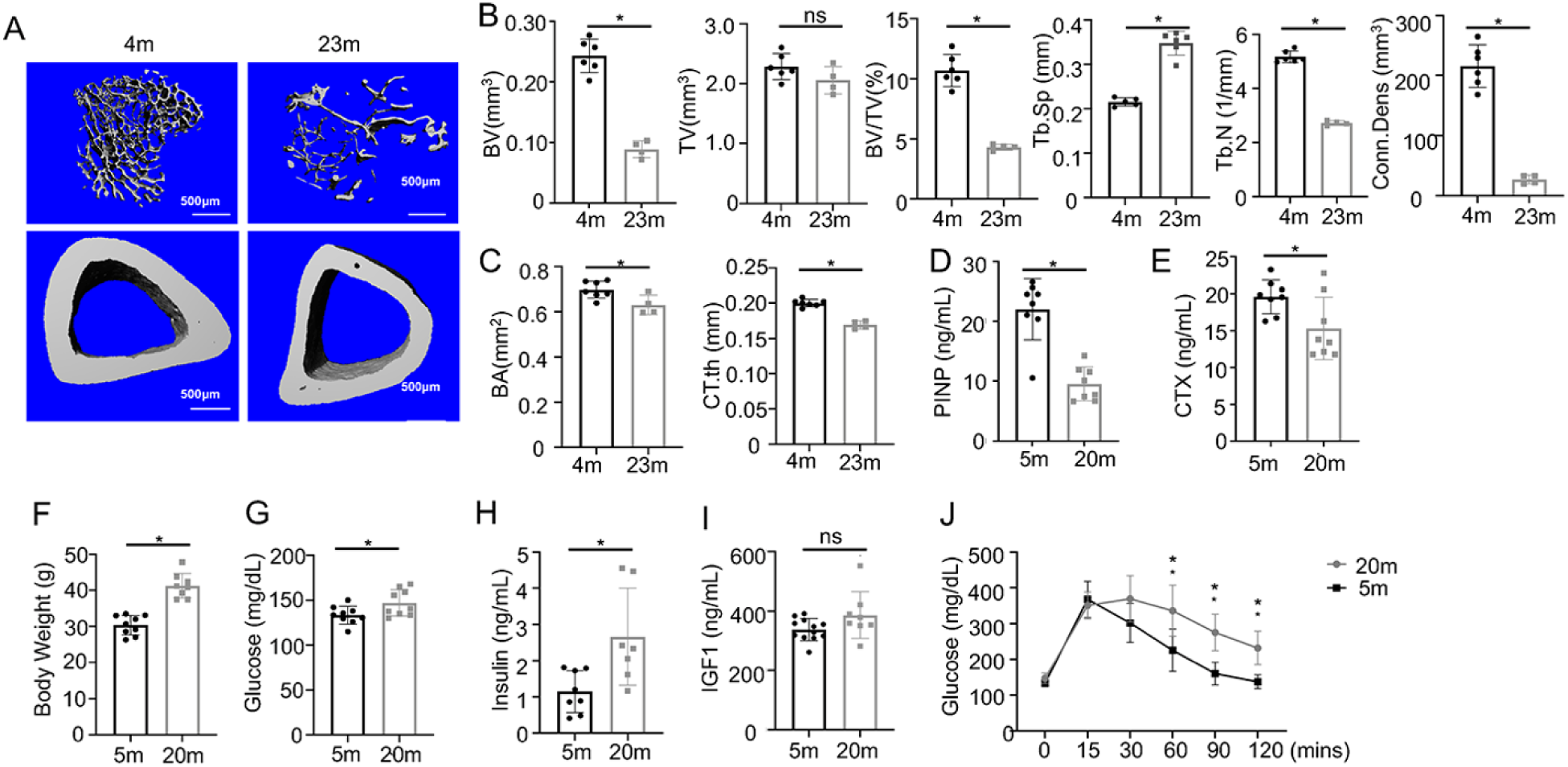
Osteoporosis is associated with impaired glucose metabolism in aged mice. (A) Representative micro-CT images of the proximal tibia (trabecular bone, upper panels) and midshaft (cortical bone, lower panels) from 4- or 23-month-old mice. (B) Quantitative analysis of trabecular bone microarchitecture in the proximal tibia, including bone volume (BV), tissue volume (TV), bone volume fraction (BV/TV), trabecular separation (Tb.Sp), trabecular number (Tb.N), and connectivity density (Conn.Dens). (C) Quantitative analysis of cortical bone parameters at the femoral midshaft, including bone area (BA) and cortical thickness (Ct.Th). (D-E) Serum levels of bone turnover markers: (D) P1NP (bone formation marker) and (E) CTX-1 (bone resorption marker). (F-I) Metabolic parameters: (F) Body weight, (G) fasting blood glucose, (H) serum insulin levels, and (I) serum IGF-1 levels. (J) Glucose tolerance test (GTT). Blood glucose was measured at indicated time points after intraperitoneal glucose injection. Data are presented as mean ± SD; *p<0.05 vs. adult group; ns, no statistically significant differences.

We next evaluated the general metabolic status of the aged mice. Aged mice displayed a significant increase in body weight (Fig. 1F). They also exhibited features of impaired glucose metabolism, including elevated fasting blood glucose and insulin levels (Fig. 1G, H), while IGF-1 protein levels remained relatively stable (Fig.1I). Moreover, glucose tolerance tests (GTT) revealed a significant impairment in glucose clearance in aged mice (Fig. 1J), which together with elevated fasting insulin suggests the presence of systemic insulin resistance. Collectively, these data support the notion that bone loss in aging is associated with a low bone turnover state and coincides with systemic metabolic dysfunction.

### Metabolomics reveals impaired glycolysis in bone

To probe potential age-related metabolic changes specifically in bone, we first performed metabolite profiling in the 20-versus 5-month-old mice. The results revealed significantly lower levels of glucose, pyruvate, lactate normalized to bone weight in the aged bone (Fig. 2A). Among the TCA cycle intermediates, α-ketoglutarate (α-KG) was significantly reduced in the aged bone whereas citrate, succinate and fumarate did not exhibit consistent changes (Fig. 2A). These findings indicate that glycolysis is likely compromised in the bone tissue of aged mice although reduced glucose uptake alone could also lead to lower levels of downstream metabolites.

**Figure 2.**
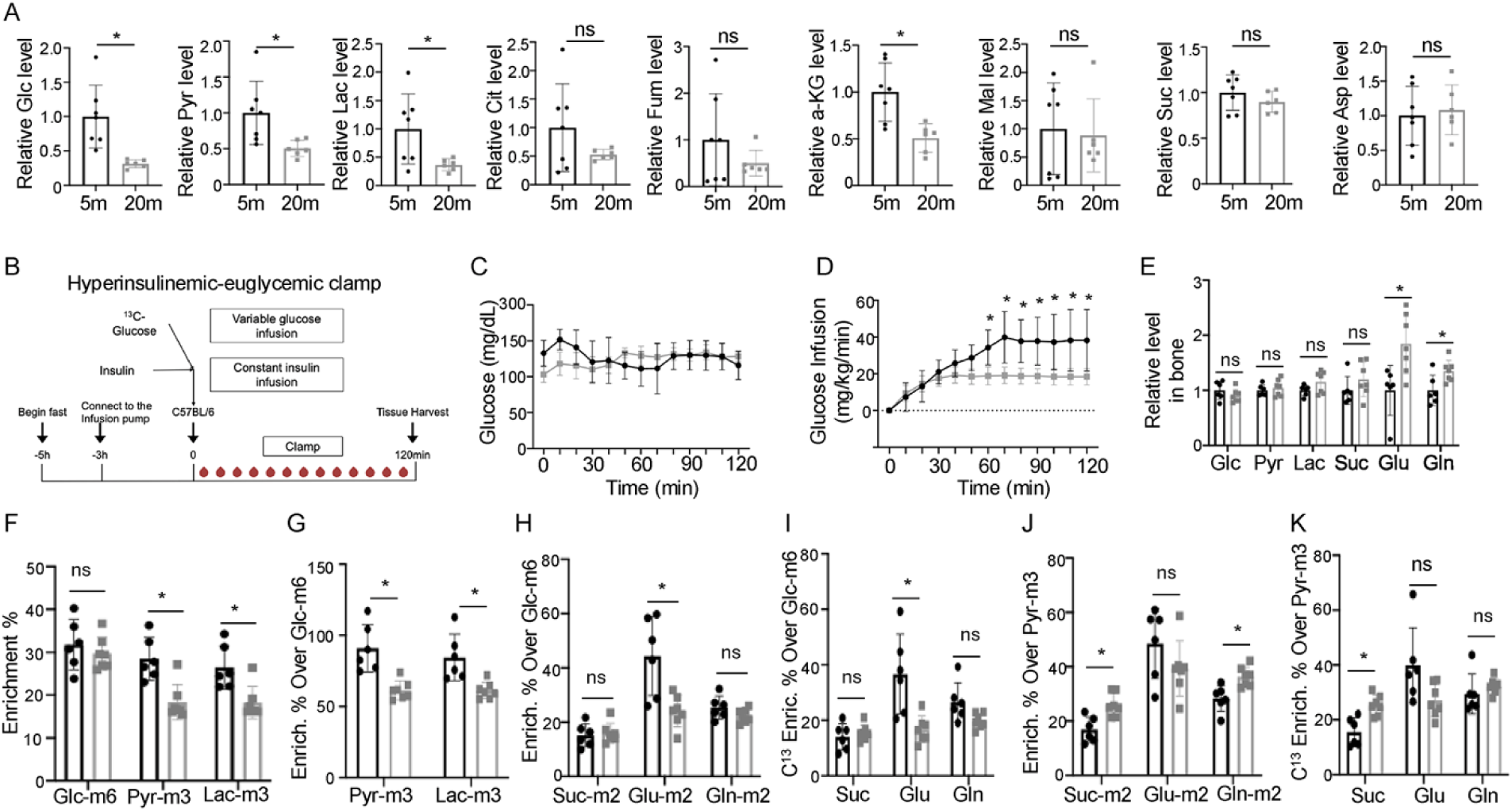
Glucose metabolism is impaired in aged bone. (A) Metabolomic analysis of bone tissue from adult and aged mice, showing steady-state levels of glucose, lactate, pyruvate, and TCA cycle intermediates (citrate, fumarate, α-ketoglutarate (α-KG), malate, succinate, and aspartate). (B) Schematic diagram of the hyperinsulinemic-euglycemic clamp procedure combined with [U-¹³C]-glucose infusion for metabolic fractional labeling analysis. (C) Blood glucose levels during the clamp procedure in adult and aged mice. (D) Glucose infusion rates required to maintain euglycemia during the clamp. (E) Absolute levels of selected metabolites in bone tissue at clamp steady state. (F) Relative abundances of ¹³C-labeled glucose, lactate, and pyruvate in bone tissue. (G) Ratios of ¹³C□-lactate/¹³C□-glucose and ¹³C□-pyruvate/¹³C□-glucose in bone tissue, indicating glycolytic fractional labeling. (H) Ratios of M+2 labeled TCA cycle metabolites to ¹³C□-glucose, representing entry of glucose-derived carbon into the TCA cycle. (I) Total carbon enrichment of TCA cycle metabolites normalized to ¹³C□-glucose. (J) Ratios of M+2 labeled TCA cycle metabolites to ¹³C_3_-pyruvate, representing entry of pyruvate-derived carbon into the TCA cycle. (K) Total carbon enrichment of TCA cycle metabolites normalized to ¹³C-pyruvate. Data are presented as mean ± SD; *p<0.05 vs. adult group; ns, no statistically significant differences.

To directly assess glucose metabolism *in vivo*, we performed a hyperinsulinemic-euglycemic clamp with [U-¹³C]-glucose infusion in young and aged mice (Fig. 2B). During the clamp procedure, blood glucose levels were successfully maintained at comparable levels of ∼150 mg/dL in both groups with continuous insulin infusion (Fig. 2C, S1A). Notably, the glucose infusion rate required to maintain euglycemia was significantly lower in the aged mice, confirming systemic insulin resistance associated with aging (Fig. 2D). Mass spectrometry analysis of metabolites in bone or plasma at the conclusion of the experiment showed that relative pool size of glucose, pyruvate and lactate was similar between the young and aged mice in both compartments (Fig. 2E, S1B). Interestingly, glutamine levels were higher in both bone and plasma of the aged mice but glutamate was higher only in bone. Additionally, citrate was increased and aspartate decreased in the plasma from aged animals (Fig. S1B). Overall, hyperinsulinemic-euglycemic clamp normalized the pool size of major glycolysis metabolites in aged bone.

We next tracked ¹³C-labeled glucose and its metabolites in bone. The relative enrichment of ¹³C□-glucose (m+6) was similar between the young and aged mice at approximately 30%; however, the enrichment of ¹³C□-pyruvate (m+3) and ¹³C□-lactate (m+3) was substantially reduced in aged bone (Fig. 2F). Likewise, the ratio of ¹³C□-pyruvate/¹³C□-glucose or ¹³C□-lactate/¹³C□-glucose was markedly lower in aged bone (Fig. 2G), indicating a bone-intrinsic reduction of glycolytic activity. Among the three TCA cycle-related metabolites detected, glutamate (m+2) (representing the first turn of the TCA cycle) or all ^13^C-labeled glutamate, normalized to glucose (m+6), was significantly reduced in aged bone, whereas the labeling of succinate or glutamine was similar between the ages (Fig. 2H, I). However, when normalized to pyruvate (m+3), glutamate (m+2) or all ^13^C-labeled glutamate no longer differed significantly between the young and aged mice, whereas succinate (m+2), glutamine (m+2) and all ^13^C-labeled succinate were higher in the aged bone, indicating that pyruvate oxidation in the TCA cycle was not impaired (Fig. 2J, K). These findings collectively indicate that impaired glycolysis is likely a root cause for bioenergetic deficiency in aged bone.

To explore whether the glycolysis defect in bone reflected global metabolic changes in aged mice, we analyzed ¹³C-labeled metabolites in the plasma. The enrichment of glucose (m+6), pyruvate (m+3) or lactate (m+3) was higher in the plasma than in bone for both young and aged mice (Supplemental Fig. S1C). Contrary to bone, the plasma glucose (m+6) enrichment was significantly lower in the aged mice, whereas pyruvate and lactate both trended lower without reaching statistical significance (Supplemental Fig. S1C). Importantly, no difference was detected in the relative enrichment of pyruvate (m+3) or lactate (m+3) normalized to glucose (m+6) in the plasma of young versus aged mice, indicating relatively normal glycolysis in the whole body (Supplemental Fig. S1D). On the other hand, the labeling fraction of TCA cycle related metabolites generally trended downward in the plasma of aged mice, with citrate and succinate reaching statistical significance (Supplemental Fig. S1E). After normalization to glucose (m+6), citrate and aspartate labeling were significantly lower in the aged plasma, suggesting an age-dependent decrease in the contribution of glucose to the TCA cycle (Supplemental Fig. S1F). Thus, the age-related changes in glycolytic metabolites in bone likely reflects tissue-intrinsic suppression of glycolysis, instead of a bystander effect of global metabolic changes.

### Transcript and protein analyses indicate dysregulation of glycolysis pathway in aged bone

The impaired glycolysis activity may reflect gene expression changes during skeletal aging. To test this possibility, we analyzed the published RNA-seq dataset from the long bone diaphysis of 3-vs 18-month-old mice (18). Principle component analyses (PCA) identified one outlier out of each age group and we therefore limited the subsequent analyses to three samples per group (Fig. 3A, Supplemental Fig. S2). GSEA analyses identified ribosome biogenesis and pathways related to this highly energy-intensive process, which is essential for protein synthesis, as the top suppressed pathways in the aged bone (Fig. 3B). Similarly, chromosome organization and several related pathways were markedly suppressed by aging (Fig. 3B). These alterations likely contributed to genomic instability and loss of proteostasis in aged bone, both considered hallmarks of aging in general (19). Targeted analyses of the differentially expressed genes detected significant alterations in glucose metabolism pathways (Fig. 3C). Interestingly, multiple core glycolysis genes including *Pfkm*, *Eno2*, *Eno3*, *Aldoa*, *Tpi1*, along with a key promoter of aerobic glycolysis *Foxk1*, were upregulated in aged bone (Fig. 3D) (20). Consistent with the RNA seq data, qPCR analysis of aged versus young bone marrow stromal cells (BMSCs) showed upregulation of *Slc2a3*, *Pfkfb3*, and *Ldha* in the aged BMSCs (Fig. 3E). In contrast, Western blot analysis revealed that the protein levels of HK2, PFKFB3 and LDHA all trended lower in aged bone, although only the decrease in PFKFB3 reached statistical significance (Fig. 3F). Thus, the increased mRNA levels of glycolysis genes likely reflect a compensatory mechanism for reduced protein levels in aged bone.

**Figure 3.**
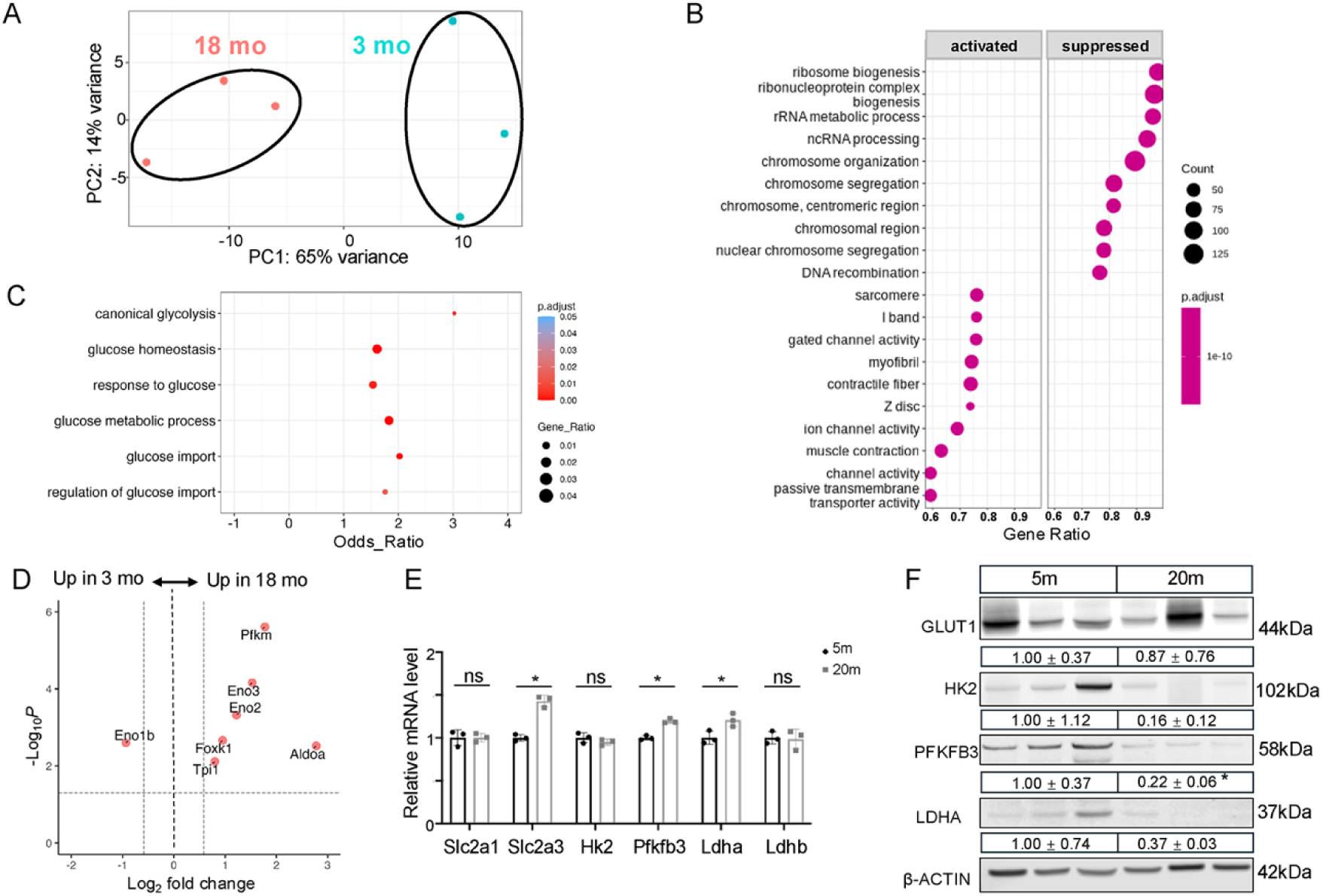
Gene expression and protein analyses indicate glycolysis changes in aged bone. (A) Principal component analysis of bulk RNA seq samples from diaphysis of 3 mo vs 18 mo mice (n=3 per age). (B) Gene set enrichment analysis depicting most highly activated or suppressed pathways in 18 mo compared to 3 mo samples. Activated means more highly active in 18 mo. (C) Dot map depicting changes in expression for glycolytic pathways. (D) Volcano plot highlighting altered canonical glycolysis pathway genes. (E) qPCR analysis of key glycolysis genes in young (5□month) versus aged (20□month) BMSCs. (F) Western blot analysis of key glycolysis enzymes in aged versus young bone tissues.

### Caloric restriction ameliorates age-related bone loss

Given the association of glycolytic impairment with age-related bone loss, we investigated whether metabolic interventions could restore bone homeostasis. Previous studies have demonstrated the beneficial effects of caloric restriction (CR) on health span and systemic glucose metabolism (21, 22). The effect of CR on bone mass, however, can be either negative or positive depending on the age of the animals (23–25). Here we analyzed mice that underwent 40% caloric restriction starting at 4 months of age and were harvested at 8 or 20 months of age. Micro-CT analysis showed that CR effectively reduced trabecular bone loss in 20-month-old mice compared to the same-age controls fed ad libitum (AL), as reflected in bone volume (BV), bone volume fraction (BV/TV) and connection density (Fig. 4A, B). In fact, the trabecular bone parameters in the 20-month-old CR mice were similar to those of the 5-month-old AL mice, indicating near complete protection from aging-associated bone loss. However, CR appeared to have no protective effects on the cortical bone, as bone area (BA) and cortical thickness (TRI-CT.th) were similarly diminished in the 20-month-old mice regardless of CR or AL (Fig. 4C). On the other hand, CR appeared to have no significant effect on the trabecular bone parameters in 8-month-old mice. This result seemed to be at odds with a previous study that CR reduced trabecular bone in mice up to one year of age, but that study used alternate-day feeding instead of the daily restricted feeding here (25). Furthermore, that study did not examine the cortical bone, but we show here that CR reduced the cortical bone area (BA) in the 8-month-old mice (Fig. 4C). The restriction of cortical bone growth by CR here mirrored a previous observation in 6-month-old mice after 3 months of CR (24). Overall, the results indicate that long-term CR protected trabecular but not cortical bone loss associated with aging.

**Figure 4.**
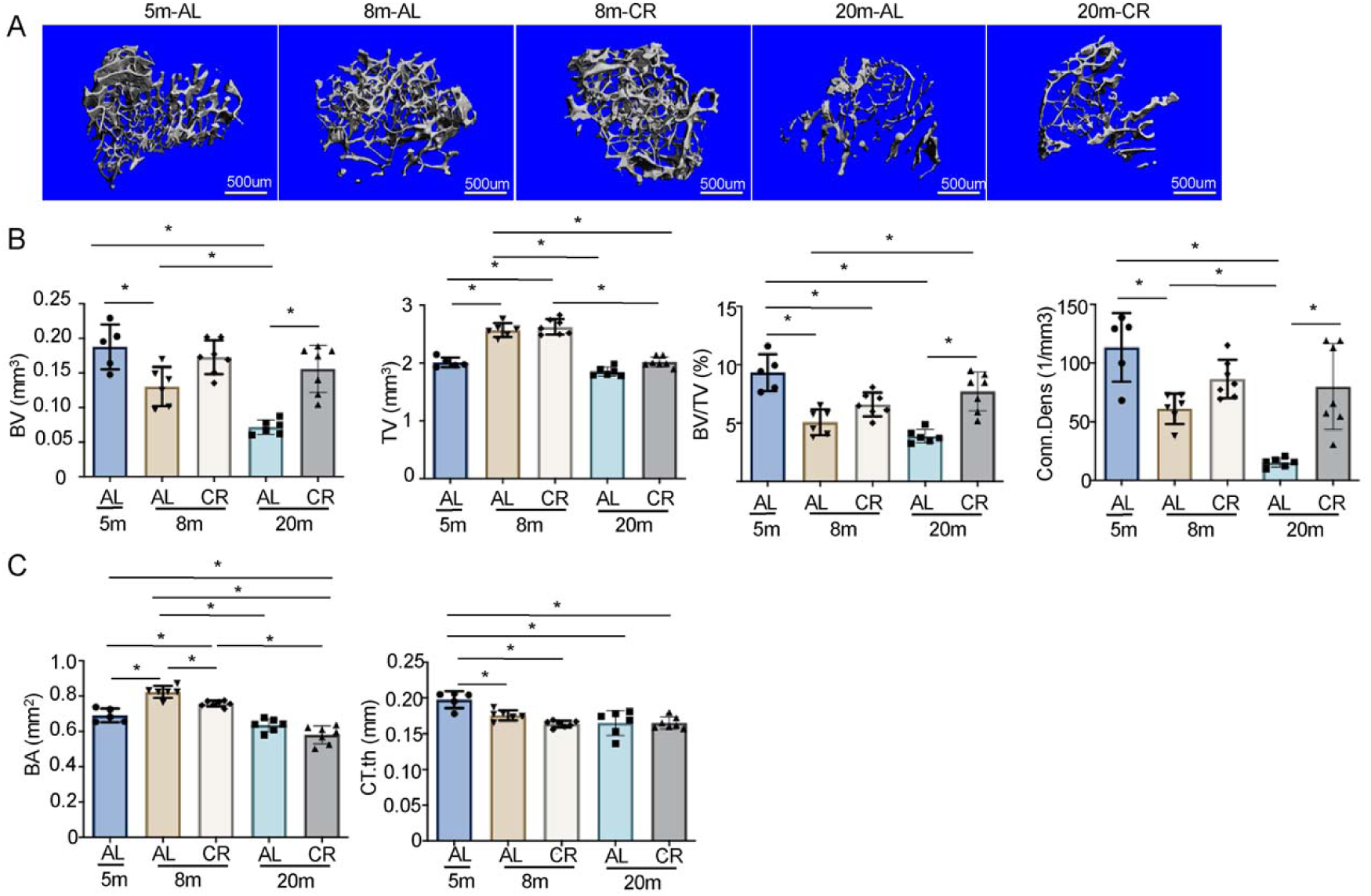
CR rescues bone loss in aged mice. (A) Representative micro-CT images of proximal tibia from 5-month-old (5M), 8-month-old ad libitum (8M-AL), 8-month-old CR (8M-CR), 20-month-old ad libitum (20M-AL), and 20-month-old CR (20M-CR) mice. (B, C) Quantitative micro-CT analysis of trabecular bone parameters (B) and cortical bone parameters (C).

### Forced expression of NAMPT does not prevent bone loss in aged mice

Since nicotinamide adenine dinucleotide (NAD^+^) is a crucial cofactor linked with glycolytic activity, we next measured total NAD (NAD^+^ plus NADH) levels in the bone tissue and found a significant decrease in aged mice (Fig. 5A). To determine whether restoring NAD levels could rescue age-related bone loss, we employed Prx1-Cre to overexpress NAMPT, the rate-limiting enzyme in the NAD salvage pathway, in the limb bud mesenchyme (26). qPCR analysis confirmed successful NAMPT overexpression in bone tissue at 3 and 5 months of age, but not at 16 months, suggesting a potential age effect on the transgene expression (Fig. 5B). Accordingly, NAD levels were significantly increased by NAMPT overexpression in the 3-or 5-but not 16-month-old mice (Fig. 5A). Micro-CT analysis, however, detected no effect on either trabecular or cortical bone by NAMPT overexpression in either males or females at any of the ages above (Fig. 5C, D, Supplemental Fig. S3). Interestingly, in 6-week-old female mice NAMPT overexpression increased trabecular bone mass, likely reflecting a transient effect on rapidly growing bones (Supplemental Fig. S3). Overall, these results suggest that NAD is not rate-limiting for sustaining bone mass in young adult mice, but cannot address the effect on aged bone due to the diminished transgene expression.

**Figure 5.**
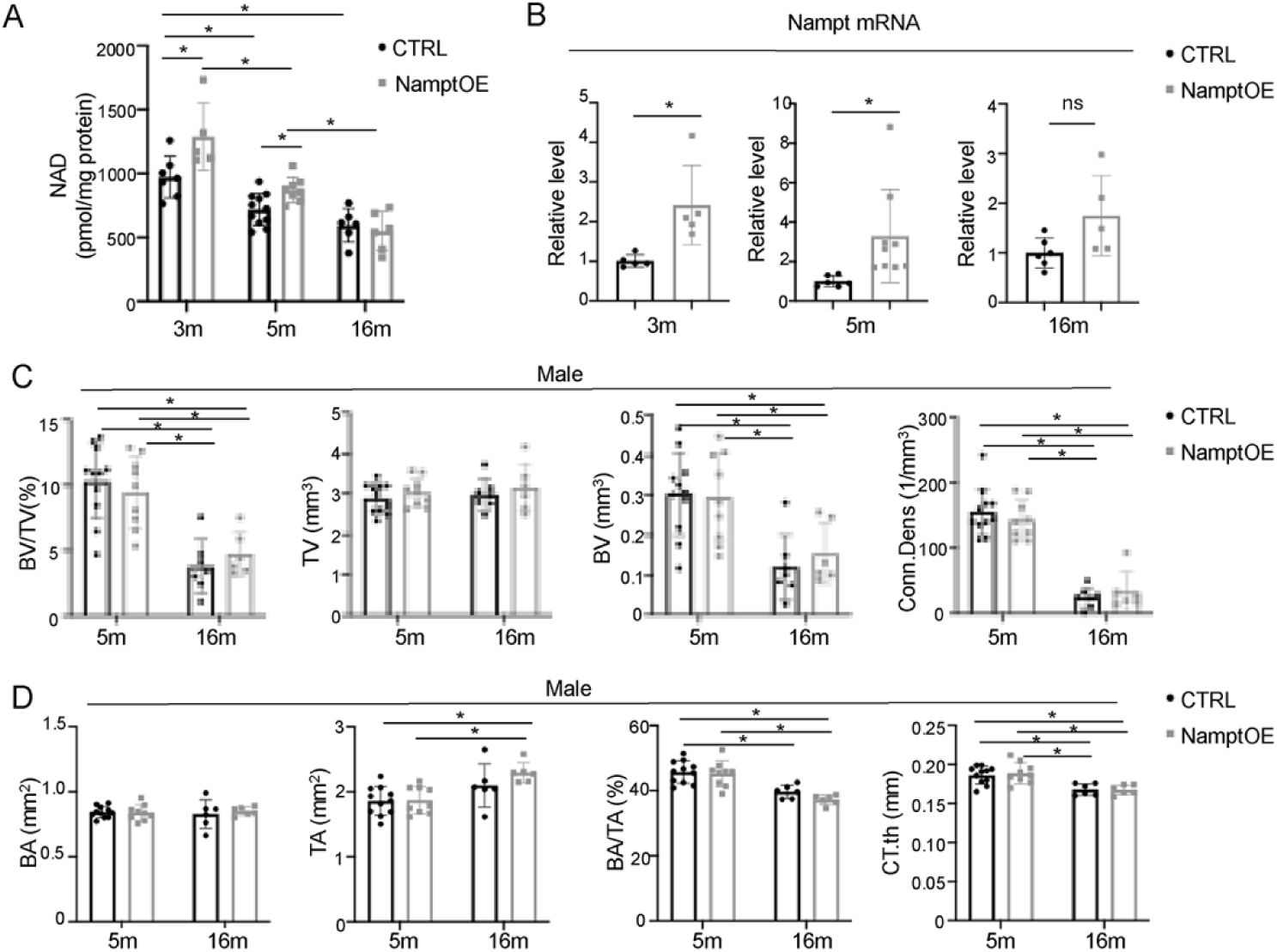
NAMPT overexpression does not improve bone loss in aged mice. (A) NAD levels in bone tissue from control and NAMPT-overexpressing mice. (B) qPCR analysis of *Nampt* expression in bone tissue from NAMPT-overexpressing mice at different ages. (C, D) Quantitative micro-CT analysis of trabecular (C) and cortical bone parameters (D) in control and NAMPT-overexpressing mice. Data are presented as mean ± SD; *p<0.05 vs age-matched controls.

### HIF1A overexpression promotes bone formation and ameliorates bone loss in aged mice

We next tested the potential effect of HIF1A, a master activator of glycolysis, on age-related bone loss. Previous work has shown that overexpression of HIF1A in osteoblasts promotes glycolysis and increases bone formation in young adult mice (27). For this, we generated mice with the genotype of Osx-rtTA;tetO-Cre;R26-Hif1a (Hif1a-OE) and induced HIF1A expression via Dox food starting at 12 months until harvest at 16 months of age. Only females were analyzed here as the males were used for another study. Micro-CT imaging showed an extended trabecular bone region in the Hif1a-OE mice (Fig. 6A). Quantification confirmed that the trabecular bone fraction (BV/TV) in the lower part of metaphysis (ROI2, 400 sections beyond ROI1) was notably increased in the Hif1a-OE mice compared to control, mainly attributable to increased trabecular thickness (Fig. 6B). The trabecular bone mass in ROI1 (400 sections beginning 100 sections below the growth plate) also appeared to increase in the Hif1a-OE mice but did not reach statistical significance (Supplemental Fig. S4A). Cortical bone parameters remained unaffected by HIF1A overexpression (Supplemental Fig. S4B). Dynamic histomorphometry in the trabecular bone showed a significant increase in bone formation rate (BFR/BS) mainly due to increased mineral apposition rate (MAR) (Fig. 6C, D). Thus, HIF1A overexpression in osteoblasts increased bone formation activity, resulting in greater trabecular bone mass in aged mice.

**Figure 6.**
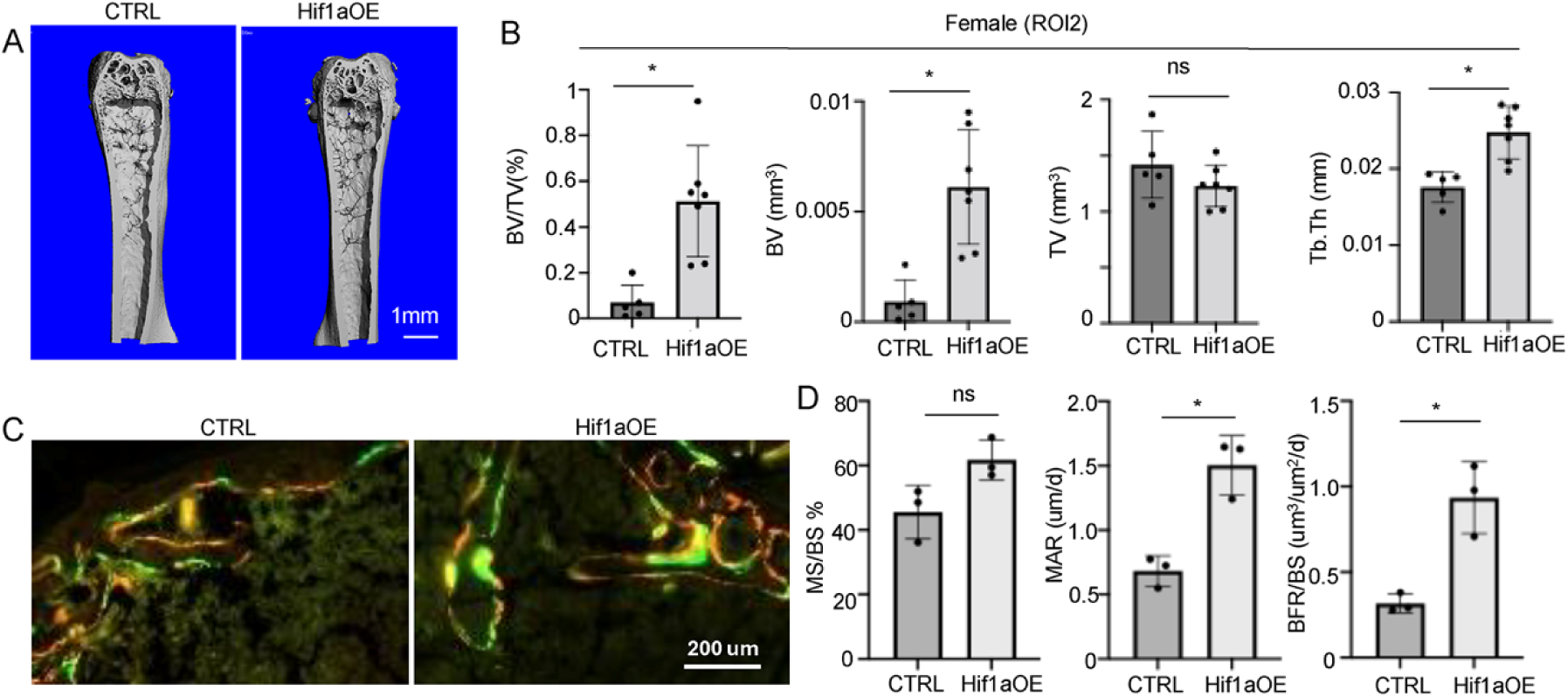
HIF1a overexpression ameliorates age-dependent bone loss. (A) Representative μCT images of the femur. (B) Quantitative analysis of trabecular bone parameters in ROI2. (C) Representative images of double labeling. (D) Quantification of double labeling. Data are presented as mean ± SD. *p < 0.05 vs. control group; ns, no statistically significant differences.

### Overexpression of PFKFB3 reduces trabecular bone loss in aged male mice

To further test the role of glycolysis in bone aging, we sought to overexpress PFKFB3 specifically in osteoblasts. PFKFB3 is transcriptionally induced by HIF1A, and produces F2,6BP that potently activates PFK-1, the first rate-limiting step of glycolysis (28, 29). We have previously demonstrated that transgenic overexpression of PFKFB3 in osteoblasts alleviated bone loss in diabetic mice (16, 17). Here, we generated mice with the genotype of Osx-rtTA;TRE-Pfkfb3 (Pfkfb3-OE) to induce PFKFB3 overexpression in osteoblasts with DOX starting at 12 months till harvest at 16 months of age.

Micro-CT analyses were performed to assess the bone parameters. 3-D reconstruction images revealed higher trabecular bone mass in the Pfkfb3-OE male mice than control (Fig. 7A). Quantification confirmed a significant increase in trabecular bone mass (BV and BV/TV), associated with increased connectivity density (Conn. Dens) and reduced trabecular separation (Tb. Sp), in ROI2 of the Pfkfb3-OE mice (Fig. 7B). ROI1 showed a trend of improvement in several parameters in the Pfkfb3-OE mice, but did not reach statistical significance (Fig. 7C). Contrary to the males, the female Pfkfb3-OE mice did not exhibit any significant changes to the trabecular bone parameters in either ROI1 or ROI2 (Supplemental Fig. S5A, B). Finally, the cortical bone parameters were not affected by PFKFB3 overexpression in either sex (Supplemental Fig. S5C, D). Thus, osteoblast-directed PFKFB3 overexpression ameliorates age-related trabecular bone loss specifically in the male mice.

**Figure 7.**
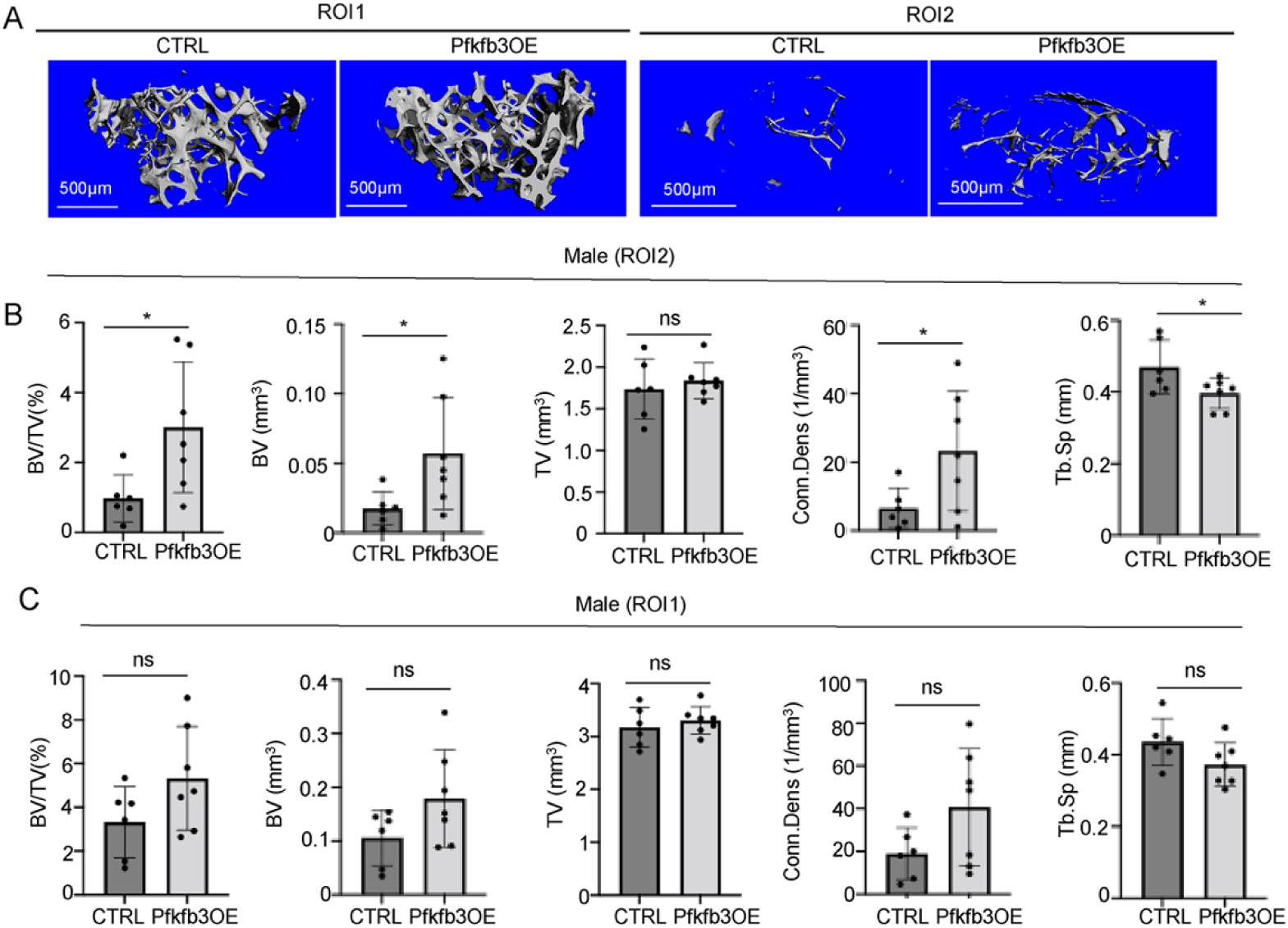
PFKFB3 overexpression reduces bone loss in aged male mice. (A) Representative μCT images of femoral trabecular bone from PFKFB3-overexpressing and control male mice. The upper panel shows images from ROI1 (a 400-slice region commencing 100 slices below the growth plate). The lower panel shows images from ROI2 (an adjacent 400-slice region directly beneath ROI1). (B, C) Quantitative μCT analysis of trabecular bone microstructure in ROI2 (B) and ROI1 (C). Data are presented as mean ± SD. *p < 0.05 vs. control group; ns, no statistically significant differences.

## Discussion

Age-related osteoporosis presents a major clinical challenge, primarily driven by a progressive decline in osteoblast activity (6). While several mechanisms have been reported, a potential role for metabolic changes in age-dependent osteoblast deficiency has not been investigated to date (5, 6). Our study identifies a notable impairment of glycolysis selectively in the bone tissue of aged mice, and further demonstrates that targeted activation of this pathway in osteoblasts can ameliorate bone loss in aged mice. The findings position the restoration of osteoblastic glycolytic activity as a potential anabolic strategy for treating age-related osteoporosis.

Glucose tracing under the conditions of hyperinsulinemic-euglycemic clamp represents a major strength of the study. This technique ensures similar blood glucose levels between the young and aged mice, minimizing the influence of variable substrate availability on downstream metabolism. The data demonstrate a clear decline in the labeling of downstream glycolytic metabolites (pyruvate and lactate) in aged bone, despite identical glucose labeling, providing strong evidence for diminished glycolysis in the bone tissue of old mice. Remarkably, this glycolysis defect was dissociated from systemic metabolism, as serum lactate or pyruvate enrichment was unaffected. Thus, aging likely impairs glycolysis in the bone tissue directly. The in vivo studies however do not distinguish specific effects on osteoblasts, osteocytes or osteoclasts.

Given the reduced glycolytic activity, the upregulation of core glycolysis genes in the transcriptomic data is unexpected. Western blot analyses, on the other hand, detected a lower level of several glycolysis enzymes in the bone tissue of aged mice. Thus, the increased mRNA transcripts may reflect a compensatory mechanism for reduced protein levels in aged bone tissues. Future work is necessary to elucidate the potential feedback mechanism.

The role of NAD^+^ in bone aging warrants further investigation. Given that NAD is a critical cofactor for glycolysis and its levels decline with age, we hypothesized that restoring NAD bioavailability might rescue bone formation in aged mice and represent a tractable therapeutic target (30–32). For this, we overexpressed NAMPT, the rate-limiting enzyme in the NAD^+^ salvage pathway, specifically in osteoblasts. The strategy modestly increased NAD levels in bone but was effective only up to 5 months of age, with no effect at 16 months. Moreover, no bone protective effect was seen in 5-month-old mice, indicating that NAD levels were unlikely to be rate-limiting for bone formation at this age. Interestingly, systemic NAD boosting in mice with nicotinamide mononucleotide starting at 5 months of age was shown by others to increase bone mineral density at 17 months of age (33). Similarly, adding nicotinamide riboside to the drinking water from 12 to 20 months of age counteracted bone loss (34). Those studies however do not distinguish systemic versus direct effects of increased NAD on bone. Future studies with strategies effectively boosting NAD levels in aged bone will clarify the potential role of bone-intrinsic NAD decline in age-dependent bone loss.

The results from HIF1A or PFKFB3 overexpression are promising. Given their role as potent activators of glycolysis and our previous characterization of the transgenic models, we expected both strategies were effective in enhancing glycolysis specifically in osteoblasts, which was most likely responsible for the bone anabolic effect (16, 17, 27). Interestingly, in both models the most notable increase in bone mass was observed in the lower half of the trabecular region. Whether or not such region-specific effect reflected varying degrees of bone formation-resorption coupling is currently unknown. Alternatively, the increase in the lower trabecular region could indicate an increased number of osteoprogenitors undergoing osteoblastogenesis in response to enhanced glycolysis. The male-specific effect of PFKFB3 overexpression is not understood at present, and could reflect varying reliance on, or regulation of, glycolysis between sexes. Regardless, sexually dimorphic response to hormonal or genetic manipulations is frequently observed in bone (35, 36). Overall, the genetic studies provide proof of principle that enhancing glycolysis is a promising bone anabolic approach for treating age-related bone loss. Future directions include identifying safe pharmacological means to stimulate the metabolic pathway selectively in osteoblasts.

## Acknowledgements

The work is partially supported by NIH grants R01DK125498 and R01AG077911 (FL), K99HL156052 and R00HL177827 (CST), R01DK098656 (JAB). The PENN Rodent Metabolic Phenotyping Core (RRID: SCR_022427) is supported in part by National Institutes of Health (NIH) grant S10-OD025098, the Cox Institute, and the Institute for Diabetes, Obesity and Metabolism (Perelman School of Medicine, University of Pennsylvania). MRM was supported by the Howard Hughes Medical Institute (HHMI) Hanna H. Gray Fellows Program Faculty Phase (Grant# GT15655) and the Burroughs Welcome Fund PDEP Transition to Faculty (Grant# 1022604).

## Author Contributions

FS, TM, WDL, JKS, CS, XL, XJ, KC, MRM conducted experiments. FS and FL designed the study and wrote the paper. JAB provided reagent. CT analyzed data. WDL, JKS, KC, MRM, JAB, CT edited the paper.

## Methods

### Sex as a biological variable

Our study examined male and female animals, and sex dimorphic effects are reported.

### Mice

All animal procedures were approved by the Institutional Animal Care and Use Committee at the Children’s Hospital of Philadelphia or University of Pennsylvania. Mice were maintained under a 12-hour light/dark cycle at 22°C. Transgenic mouse lines including TRE-PFKFB3, Osx-rtTA, Hif1dPA, and TetO-Cre, were as previously described and maintained on a C57BL/6J background (16, 37–39). For inducible expression, doxycycline (Dox; 1 mg/mL, D9891, Sigma) was administered in drinking water. For the ^13^C-glc hyperinsulinemic-euglycemic clamp experiment, young and aged C57BL/6J male mice were purchased from the Jackson Laboratory. For CR studies, C57BL/6J.Nia male mice either ad libitum fed (AL) or 40% caloric restricted (CR, starting at 4 months of age) were obtained from National Institute on Aging Rodent Colony and maintained on NIH-31 and NIH-31 fortified diet, respectively, for two months prior to use, during which time CR mice were single housed and fed a 3g pellet everyday between 8–10 am.

### Skeletal phenotyping

The proximal tibiae of young (5-month-old) and aged (20-month-old) male mice were scanned using a μCT35 system (SCANCO Medical) at an isotropic voxel size of 7 µm. A stack of 200 slices was analyzed per sample for trabecular bone quantification. The distal femur of each animal was scanned using a μCT45 system (SCANCO Medical) at an isotropic voxel size of 4.5 µm. For trabecular bone analysis, two non-overlapping regions of interest (ROI1 and ROI2) were defined in the distal femur metaphysis: ROI1 extended from 0.45 mm to 2.25 mm, and ROI2 extended from 2.25 mm to 4.05 mm below the growth plate. For cortical bone analysis, a 70-slice region centered at the femoral mid-diaphysis, located 5.4 mm from the distal growth plate, was evaluated. The following parameters were calculated: for trabecular bone—bone volume (BV), tissue volume (TV), bone volume fraction (BV/TV), trabecular thickness (Tb.Th), trabecular number (Tb.N), trabecular separation (Tb.Sp), and connectivity density (Conn.Dens); for cortical bone—total area (TA), bone area (BA), bone area fraction (BA/TA), and cortical thickness (Ct.Th).

Dynamic histomorphometry was performed by administering sequential intraperitoneal injections of calcein green (5 mg/kg) and alizarin red (15 mg/kg) at 7 days and 2 days, respectively, prior to euthanasia. Following dissection, femurs or tibias were fixed in 4% PFA in PBS for 48 hours, then stored in 70% ethanol in the dark at 4°C until further processing. Prior to sectioning, samples were cryoprotected in 30% sucrose in PBS overnight and embedded in OCT compound. Undecalcified longitudinal sections (10 µm thick) were prepared using a cryostat with Cryofilm type II membrane (Section Lab, Co. Ltd., Japan). Fluorescence imaging was carried out on an Axioscan slide scanner (ZEISS, Germany), and subsequent quantification was performed using the BIOQUANT Image Analysis System (BIOQUANT Image Analysis Corporation, USA).

### Serum biochemical assays

Following a 6-hour fasting period, blood samples were collected from mice via retro-orbital bleeding. Serum levels of CTX-I and P1NP were measured using the RatLaps (CTX-I) EIA and Rat/Mouse P1NP EIA Kit (Immunodiagnostic Systems, Ltd.), respectively. Serum insulin concentration was determined with an ultra-sensitive mouse insulin ELISA kit (Crystal Chem, Cat#90080). Fasting blood glucose was measured directly from a tail-vein sample using a standard glucometer.

### Relative quantification of metabolites *in vivo*

Young (5-month-old) and aged (20-month-old) C57BL/6J mice were acclimatized to the animal facility for one week under a 12-hour light/dark cycle (7:00 AM to 7:00 PM) with ad libitum access to food and water. A jugular vein catheter was surgically implanted, and hyperinsulinemic-euglycemic clamps were performed within one week post-surgery in the PENN Rodent Metabolic Phenotyping Core. A variable infusion of [U-¹³C□]-glucose and a constant infusion of insulin (2.5 mU/kg/min) were administered to achieve and maintain steady-state blood glucose at approximately 150 mg/dL for 2 hours. Mice were connected to a tether and swivel system (Instech Laboratories) to allow free movement with continuous access to food and hydrogel (Clear H□O) throughout the infusion. Following the clamp, plasma and bone shaft samples were collected.

The bone shaft of long bones was dissected quickly, with the metaphysis excised and bone marrow removed by centrifugation. The bone shaft was snap frozen in liquid nitrogen and stored at –80□°C until metabolite extraction. At that time, the frozen bone tissue was weighed into a pre□chilled tube and ground under liquid nitrogen using a cryomill (Retsch) at 25□Hz for 45□seconds. To the resulting bone powder, –80□°C 80:20 methanol:water was added at a volume of 40□μL solvent per 1□mg of tissue. The mixture was vortexed for 15□seconds, incubated on dry ice for 10□minutes, and then centrifuged at 16,000□g for 20□minutes at 4□°C. The supernatant was transferred to a new Eppendorf tube and centrifuged again at 16,000□g for 25□minutes to remove any residual debris. The final supernatant was used for LC MS analysis as previously described (17). Metabolites were analyzed using a Vanquish Horizon UHPLC System (Thermo Scientific) coupled to an Orbitrap Exploris 480 Mass Spectrometer (Thermo Scientific). Waters XBridge BEH Amide XP Column (particle size, 2.5□μm; 150□mm (length)□×□2.1□mm (i.d.)) was used for hydrophilic interaction chromatography (HILIC) separation. Column temperature was 25 °C. Mobile phases A□=□20 mM ammonium acetate and 22.5 mM ammonium hydroxide in 95:5 (v/v) water:acetonitrile (pH 9.45) and B□=□100% acetonitrile were used for both ESI positive and negative modes. The linear gradient eluted from 90% B (0.0–2.0 min), 90% B to 75% B (2.0–3.0 min), 75% B (3.0–7.0 min), 75% B to 70% B (7.0–8.0 min), 70% B (8.0–9.0 min), 70% B to 50% B (9.0–10.0 min), 50% B (10.0–12.0 min), 50% B to 25% B (12.0–13.0 min), 25% B (13.0–14.0 min), 25% B to 0.5% B (14.0–16.0 min), 0.5% B (16.0–20.5 min), 90% B (20.5 – 25.0 min). The flow rate was 0.15 mL/min. The sample injection volume was 5 μL. ESI source parameters were as follows: spray voltage, 3200 V or −2800 V, in positive or negative modes, respectively; sheath gas, 35□arb (arbitrary units); aux gas, 10□arb; sweep gas, 0.5□arb; ion transfer tube temperature, 300 °C; vaporizer temperature, 35 °C. LC–MS data acquisition was operated under full scan polarity switching mode for all samples. The full scan was set as: orbitrap resolution, 120,000 at m/z 200; AGC target, 1e7; maximum injection time, 200 ms; scan range, 60–1000 m/z. LC-MS raw data files (.raw) were converted to mzXML format using ProteoWizard (version 3.0.20315). El-MAVEN (version 0.12.0) was used to generate a peak table containing m/z, retention time, and intensity for the peaks. For tracer experiments, isotope labeling was corrected for ^13^C natural abundances using AccuCor package. ¹³C enrichment was calculated as: Carbon Enrichment = (m□×0 + m□×1 + m□×2 + … + m□×n) / n where m□ represents the mass isotopologue fraction containing *x* ¹³C atoms, and *n* is the total number of carbon atoms in the molecule.

### NAD measurement

The bone shafts of long bones (femurs and tibias) were rapidly dissected, with the metaphyseal regions removed and bone marrow eliminated by centrifugation. The remaining cortical bone shafts were immediately snap-frozen in liquid nitrogen and stored at −80 °C until analysis. For metabolite extraction, frozen bone samples were homogenized in NAD/NADH extraction buffer provided in the NAD/NADH Assay Kit (Colorimetric) (ab65348, Abcam, Cambridge, UK). Samples were subsequently deproteinized using the supplied spin columns according to the manufacturer’s instructions. Total NAD (NADt; NAD□+ NADH) was measured directly from the deproteinized extracts. For NADH determination, aliquots of the extracts were incubated at 60 °C for 30 min to selectively decompose NAD□ before cooling on ice. Samples and standards were then loaded into 96-well plates, followed by addition of the reaction mix to convert NAD to NADH. After a 5-min incubation at room temperature, NADH developer was added to initiate the enzymatic cycling reaction, and absorbance at 450 nm was monitored using a microplate reader during the 1–4 h incubation period. NAD concentrations were calculated by subtracting NADH values from total NAD levels. The assay specifically detected NAD□ and NADH but not NADP□ or NADPH.

### Transcriptomics analysis

Raw RNA sequencing data were obtained from the National Genomics Data Center (https://ngdc.cncb.ac.cn), processed using the Nextflow nf-core RNA sequencing pipeline (Nextflow v24.04.3, nf-core/rnaseq v3.14.0), and analyzed using publicly available software platforms (DESeq2, clusterProfiler) in R (v4.2) (PMID 28398311, PMID 32055031). Coding scripts for analyzing processed data are available on Github (https://github.com/thomchr/BoneAgeMetabolism).

### Western Blot

The bone shaft was collected as described above, snap frozen in liquid nitrogen and stored at – 80□°C until Western blot analysis. Frozen bone tissue was ground under dry ice using M□PER extraction reagent (cat#78501, Thermo Scientific, Waltham, MA, USA) with a Precellys Evolution homogenizer (Bertin, Montigny□le□Bretonneux, France) at 6000□rpm for 15□seconds at 4□°C for 3 cycles. The lysate was vortexed and then centrifuged at 16,000□g for 20□minutes at 4□°C. The supernatant was collected, and protein concentration was determined using the BCA assay (Pierce™ BCA Protein Assay Kit, Thermo Fisher Scientific) with bovine serum albumin (BSA) as the standard. Equal amounts of protein were separated by SDS□polyacrylamide gel electrophoresis (SDS PAGE) and transferred onto polyvinylidene difluoride (PVDF) membranes (Millipore). Membranes were blocked with 5% non□fat dry milk in Tris□buffered saline containing 0.1% Tween 20 (TBST) for 1□hour at room temperature, followed by overnight incubation at 4□°C with primary antibodies. Alternatively, membranes were blocked with LI□COR blocking buffer (cat#927□60001) prior to primary antibody incubation. The following primary antibodies were used: anti□Glut1 (Abcam, ab1157730), anti Pfkfb3 (Abcam, ab181861), anti□Ldha (Cell Signaling Technology, 2012), anti□Hk2 (Abcam, ab228819), and anti□β□actin (Cell Signaling Technology, 3700). After washing, membranes were incubated with secondary antibodies: anti□rabbit IgG (cat#926□32211, LI□COR) or anti mouse IgG (cat#926 68070, LI□COR). Images were captured and quantified using LI□COR imaging software (LI□COR, Nebraska, USA).

### BMSCs isolation for qPCR

Bone marrow cells were flushed from tibias and femurs (epiphyses removed) and cultured in MEM α with 10% FBS. After reaching confluence (day 6–8), cells were dissociated with collagenase II and trypsin EDTA. CD45+ hematopoietic cells were depleted using magnetic activated cell sorting (MACS) with CD45 MicroBeads (Miltenyi Biotec, USA). Purified BMSCs were passaged once, then harvested with collagenase I and trypsin for RNA extraction and qPCR analysis. Relative changes were calculated as fold change using the 2□ΔΔCt method. The primer sequences are listed in Supplementary Table 1.

### Statistics

Statistical analysis was performed using GraphPad Prism software version 9.0. Differences between two groups were evaluated by unpaired t-test. For comparisons among more than two groups, one-way or two-way ANOVA was applied as appropriate, followed by post hoc testing using Sidak’s correction or Fisher’s LSD test. Data are presented as mean ± standard deviation (SD). A P value < 0.05 was considered statistically significant.

## Data availability

Analytic code for analyzing RNA-seq data is available on Github (https://github.com/thomchr/BoneAgeMetabolism).

