## Supplemental Figures and Legends for "Impaired glycolysis in bone underlies age-related bone loss in mice"

**Supplementary Figure S1. Metabolic Analysis During Hyperinsulinemic-Euglycemic Clamp, Related to Figure 2**.

(A) Plasma insulin concentrations during the clamp procedure. (B) Relative plasma levels of key metabolites. (C) Plasma enrichment of ^13^C_6_-glucose and its metabolites ^13^C_3_-lactate and ^13^C_3_-pyruvate. (D) Ratios of ^13^C_3_-lactate/^13^C_6_-glucose and ^13^C_3_-pyruvate/^13^C_6_-glucose. (E) Plasma ^13^C enrichment of TCA cycle intermediates. (F) Plasma enrichment of TCA cycle intermediates normalized to ^13^C_6_-glucose. (G) Plasma enrichment of TCA cycle intermediates normalized to ^13^C_3_-pyruvate. Data are presented as mean ± SD. *p < 0.05, Student’s t test, between age groups; ns, no statistical significance.

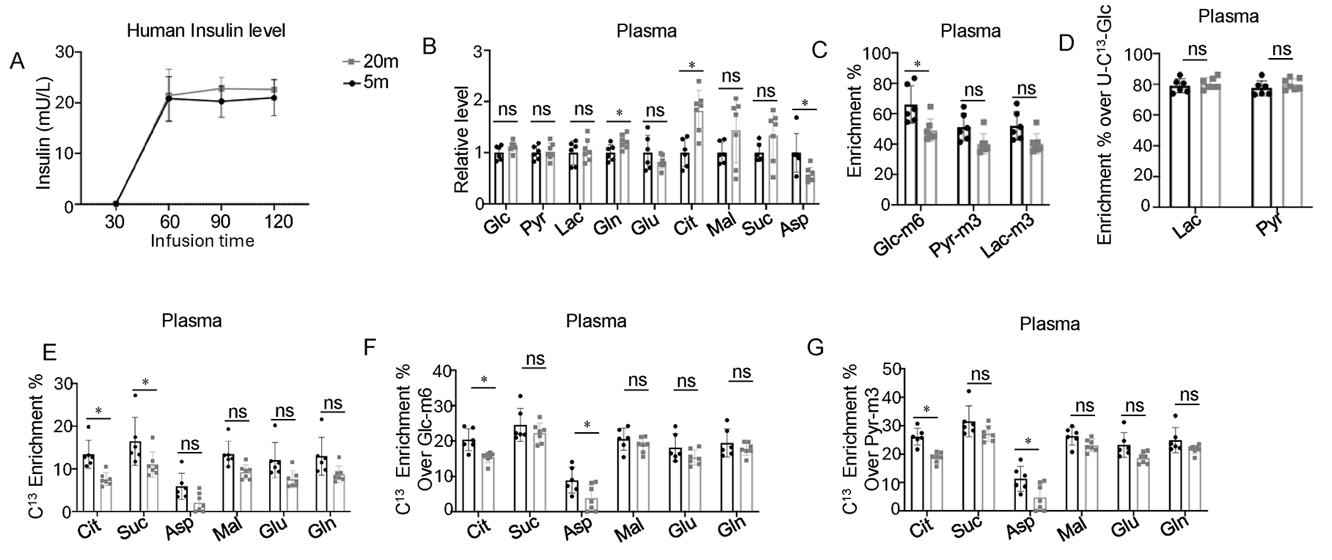

**Supplemental Fig. S2. PCA analyses of bone RNA-seq dataset. Related to Figure 3.**

Oval encircles one outlier from each age group excluded from downstream analyses.

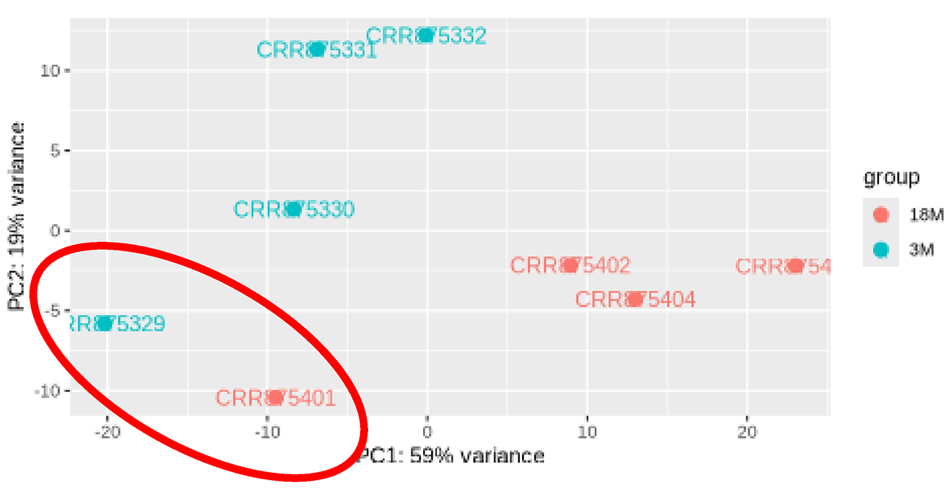

**Supplementary Figure S3. Minimal Impact of Nampt Overexpression on Bone Mass. Related to Figure 5.**

(A) µCT quantification of trabecular bone parameters in male control (black) or Nampt-OE (grey) mice. (B, C) µCT quantification of cortical (B) and trabecular (C) bone parameters in female control (black) or Nampt-OE (grey) mice. Data are presented as mean ± SD. *p < 0.05, Student’s t test between genotypes; ns, no statistically significant differences.

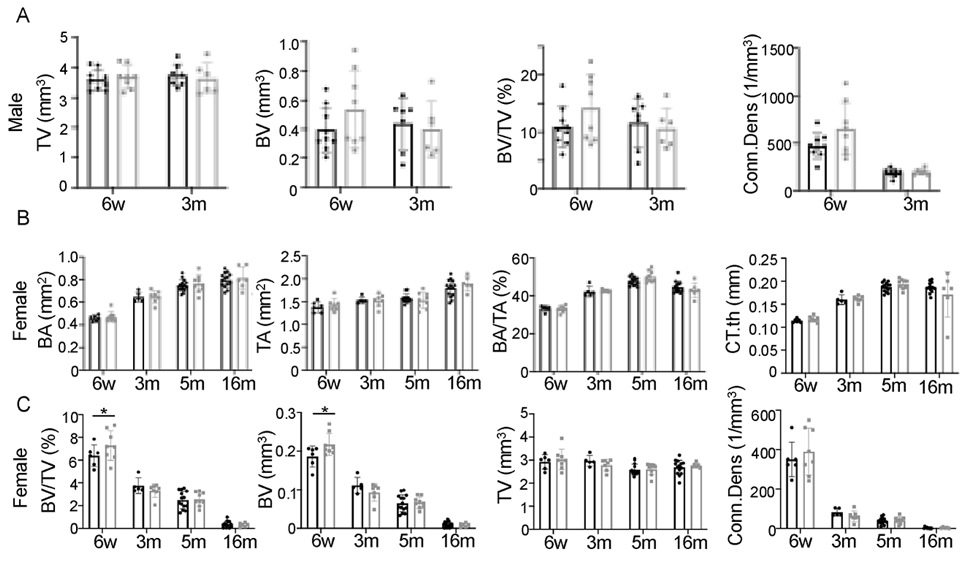

**Supplementary Figure S4. Effects of HIF1a Overexpression on Bone Microarchitecture in Aged Female Mice, Related to Figure 6.**

(A) Quantitative μCT analysis of trabecular bone in the distal femur ROI1 (a 400-slice region commencing 100 slices below the growth plate) of 16-month-old female mice. (B) Quantitative analysis of cortical bone parameters in 16-month-old female mice. Data are presented as mean ± SD. No statistical difference between groups.

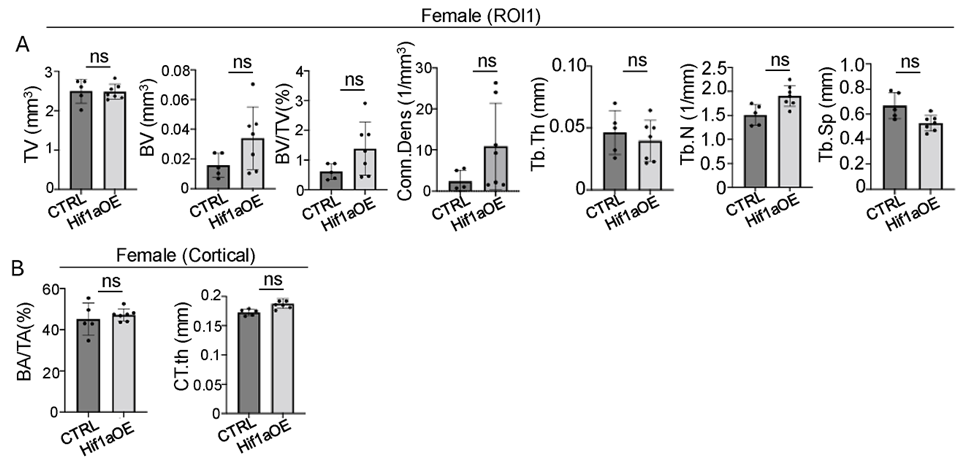

**Supplementary Figure S5. Effects of PFKFB3 Overexpression on Cortical and Trabecular Bone in Aged Mice, Related to Figure 7.**

(A, B) Quantitative μCT analysis of trabecular bone in ROI1 (A) or ROI2 (B) of the distal femur of female mice at 16 months of age. (C) Quantitative μCT analysis of femoral cortical bone in 16-month-old male (C) or female (D) mice. Data are presented as mean ± SD. No statistically significant difference between groups.

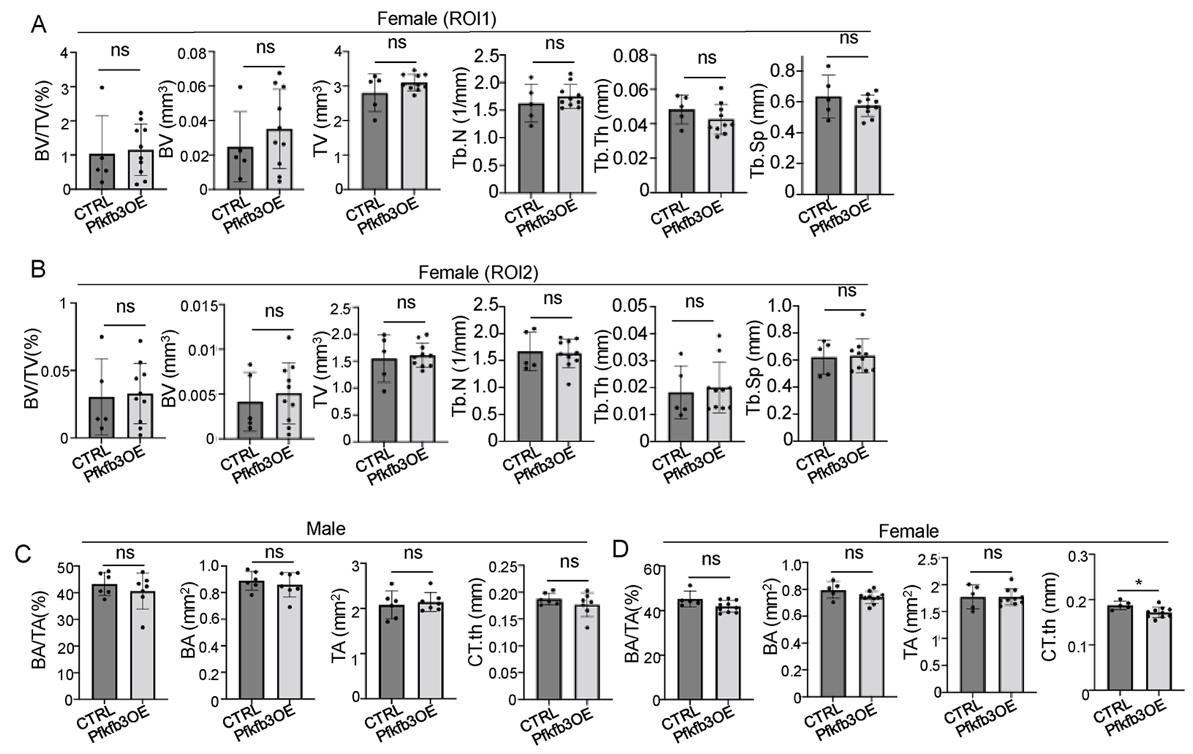
